# BICORN: An R package for integrative inference of *de novo* cis-regulatory modules

**DOI:** 10.1101/560557

**Authors:** Xi Chen

## Abstract

BICORN is an R package developed to integrate prior transcription factor binding information and gene expression data for cis-regulatory module (CRM) inference. BICORN searches for a list of candidate CRMs from binary bindings on potential target genes. Applying Gibbs sampling, BICORN samples CRMs for each gene using the fitting performance of transcription factor activities and regulation strengths of TFs in each CRM on gene expression. Consequently, sparse regulatory networks are inferred as functional CRMs regulating target genes. The BICORN package is implemented in R and is available at https://cran.r-project.org/web/packages/BICORN/index.html.

## 1 Introduction

Transcription factor (TF) binding data are widely available given the rapid development of epigenetic biotechnologies (e.g., ChIP-chip, ChIP-seq, DNase-seq and ATAC-seq), efficient genome-wide binding motif searching algorithms and large-scale databases (e.g., JASPAR, ENCODE and Roadmap). It is well known that gene transcription is mediated by cis-regulatory modules (CRMs) (1). Inferring TF associations within CRMs is essential to understand how genes are regulated by the combined effect of TFs. Several software packages can infer *de novo* CRMs using binding data (2,3). However, a major limitation of these packages from the application perspective is that they do not integrate with gene expression data and cannot tell which module is functionally regulating which target genes. Conventional TF-gene regulatory network inference tools (4–7) cannot fully solve this problem because they only predict individual TF-gene interactions.

Here, we present an R package (BICORN) to perform *de novo* functional CRM inference. Using prior TF-gene binding knowledge and gene expression data as input, BICORN infers a list of functional CRMs and their target genes. Key functionalities of BICORN include: (1) candidate CRM searching; (2) Gibbs sampling-based estimation of TF activity (TFA) and regulation strength (RS); (3) CRM sampling according to the linear fitting performance of TFAs and their regulation strengths on target gene expression.

## 2 Model Description

A workflow of BICORN is shown in Fig. 1. The BICORN software package requires two inputs (i.e., prior bindings and gene expression data) to infer CRMs. Binary prior binding data are used because it is the most common signal format from different resources. For example, many databases can provide potential target genes for a TF query, a binary binding relationship. Using ChIP-seq data, target genes can be directly inferred based on TF binding signals and gene annotation files (8,9). For DNase-seq or ATAC-seq data, binding site scanning is a straightforward approach that scans a specific TF binding motif within open chromatin regions near target genes (10). For gene expression, RNA-seq data are preferred and measured by log-transformed transcripts per million (TPM) values (11). Microarray data also can be used after proper normalization. BICORN automatically filters out genes either regulated by less than two TFs (the minimum number of TFs required in a CRM) or missing expression data. Instead of performing an exhaustive search on TF combinations (e.g., 2^50^ given 50 TFs), a list of candidate CRMs is initially identified by BICORN based on the prior binding events.

**Fig 1.**
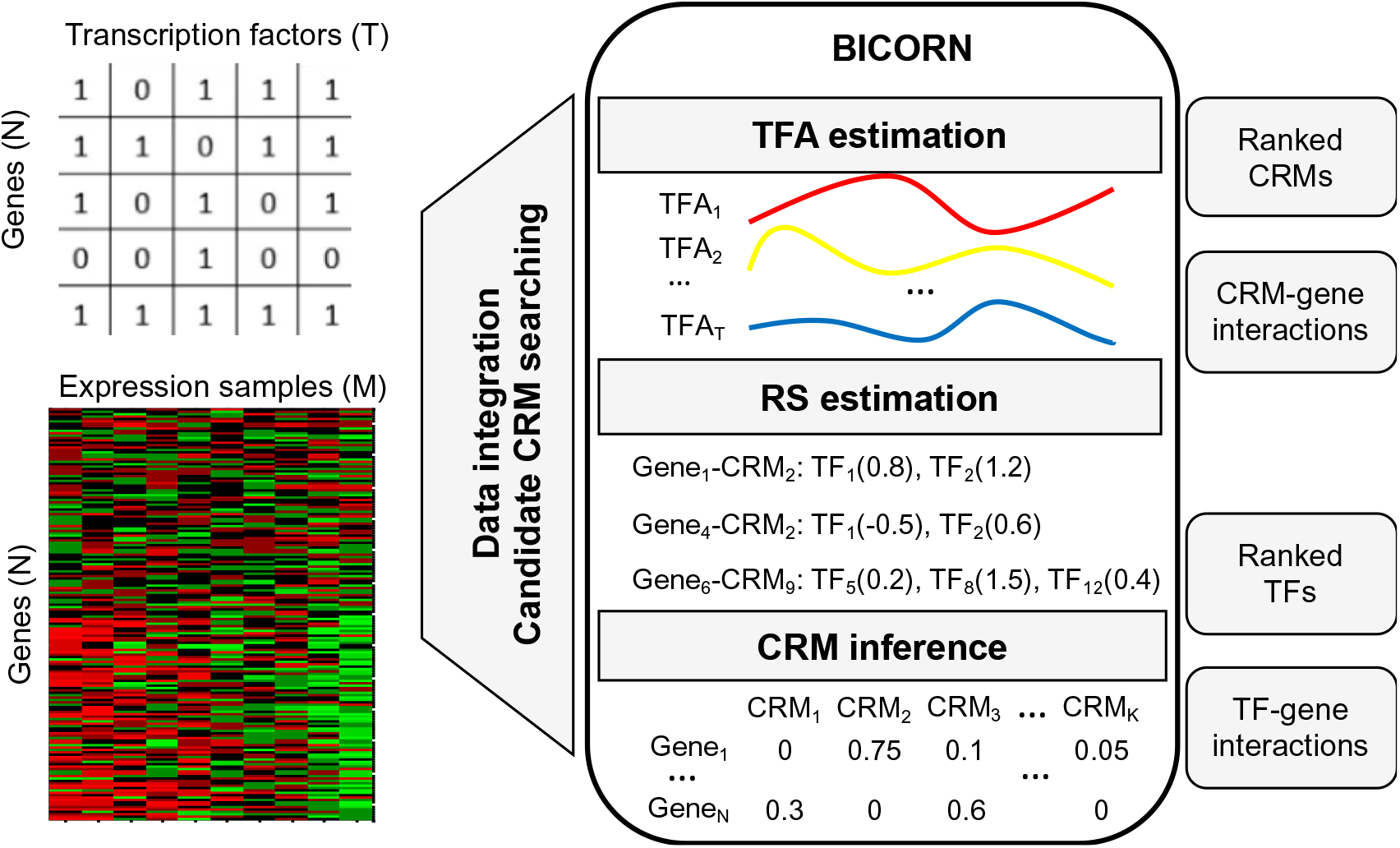
A workflow of the BICORN package.

Using a log-linear model (*Supplementary Material S1, Eq. (S-1)*), the log-transformed expression value of a gene is modeled as a linear combination of TFA, RS, and CRM variables. The CRM variable is the key variable controlling how TFs are grouped such that the association of TFs in a CRM can be directly inferred. All variables are iteratively estimated using Gibbs sampling according to their conditional probabilities. More details can be found in *Supplementary Material S1 and Fig. S1*.

### 2.1 Transcription factor activity (TFA) estimation

A key factor of BICORN that differentiates it from some existing methods is estimating TFA, which is the active proteomic response controlling target gene expression. The correlation of transcripts and proteins (especially protein activities) varies greatly depending on the cellular locations and biological functions of genes (12). In this study, a TF is treated as a regulatory protein so that it is reasonable to estimate and use TF activity rather than its mRNA expression to fit target gene expression, although the latter usually simplifies the computational complexity. Here, TFA is modeled as a Gaussian process, the value of which is sampled according to a conditional Gaussian distribution.

### 2.2 Regulation strength (RS) estimation

Given a number of TFs, most existing tools only report strong regulators for each gene (13), to lower the impact of overfitting on regulatory network inference, leading to an incomplete reconstruction of the true regulatory network. BICORN samples CRMs individually. The regulation strength (RS) of TFs in each module is jointly estimated, including both strong and weak regulators. Since the number of TFs is well controlled (between two and six) within each candidate CRM, the overfitting effect can be largely alleviated. Candidate CRMs are sampled sufficiently to ensure that all observed TF binding events for each gene are fully examined.

BICORN is especially useful when a relatively large number of TFs (*e.g.*, over 50) are jointly studied because genes can be frequently regulated by a number of TFs (e.g., 10 - 20) in the prior binding network. Thus, each gene can be potentially regulated by several CRMs. At each iteration BICORN samples only one CRM for each gene so that the gene expression level is fitted using TFAs and RSs of a small number of TFs. Compared to conventional regulatory network inference methods fitting gene expression using all the TFs simultaneously, the overfitting effect in BICORN is much lower.

### 2.3 cis-regulatory module (CRM) inference

For each target gene, at each iteration BICORN samples a CRM by applying a conditional probability of how well the gene expression can be fitted using estimated TFAs and RSs of TFs in the sampled CRM. After sufficient rounds of samplings, the sampling frequency of each CRM-gene represents the posterior probability that each regulatory event occurred. Outputs of BICORN consist a list of CRMs and a weighted CRM-gene regulatory network. BICORN also outputs a weighted TF-gene regulatory network based on inferred CRM-gene interactions.

Using simulated data, we compared the performance of BICORN with that of several existing software packages (*Supplementary Material S2, Figs. S2-S3, Table S1*). Results showed that BICORN can identify CRMs and their target genes with a higher accuracy. We applied BICORN to breast cancer MCF-7 cell-specific data and inferred CRMs functional at enhancer and promoter regions, respectively (*Supplementary Material S3, Figs. S4-S5*). As shown in Fig. S6, given the same set of TFs, 11 are active at enhancer regions while 13 are active at promoter regions, with only four TFs overlapped. Thus, TFs form distinct CRMs at enhancer or promoter regions to mediate gene expression. We finally applied BICORN to datasets of multiple cell types and demonstrated its usefulness in identifying biologically meaningful CRMs (*Supplementary Material S4, Table S2*).

