## Supplemental material for "BICORN: An R package for integrative inference of *de novo* cis-regulatory modules"

Xi Chen, Jinghua Gu, Leena Hilakivi-Clarke, Robert Clarke, Tian-Li Wang, Jianhua Xuan

### **S1 BICORN algorithm**

Key functional modules in BICORN package are illustrated in Fig. S1. It requires two inputs as prior binding knowledge and gene expression data. After data preprocessing, BICORN mainly performs three computational functions: (1) sampling of transcription factor activity, (2) sampling of regulation strength and (3) sampling of cis-regulatory modules (CRMs) for *de novo* CRM inference.

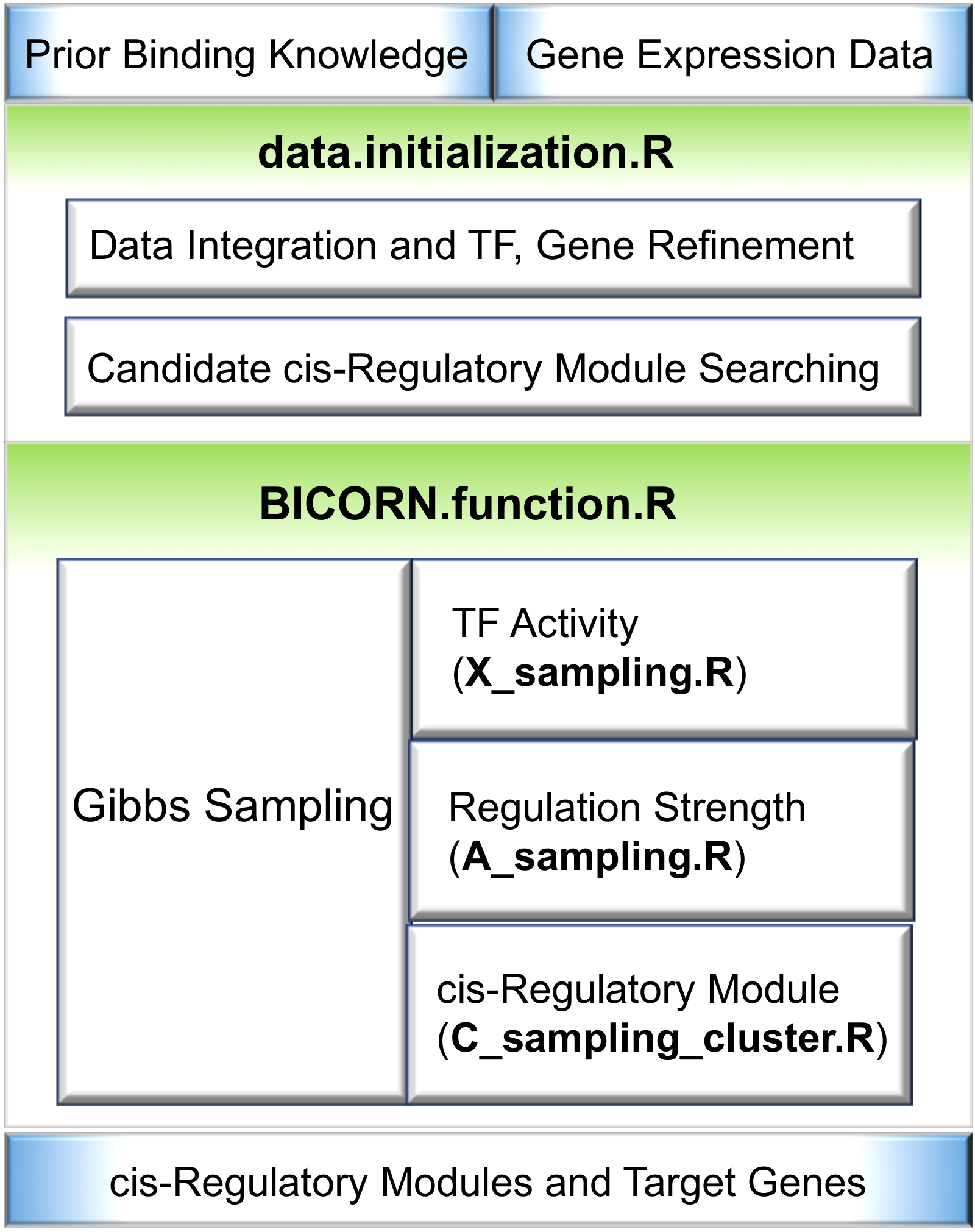

**Figure S1**. Key functions of the BICORN Package.

#### **S1.1 Prior binding knowledge and candidate CRM searching**

Given the prior binding knowledge and gene expression, BICORN infers the most reliable cis-regulatory module (CRM) for each gene. Prior binding knowledge can come from different data recourses and databases. What we need is the binary binding relationship between transcription factors (TFs) and potential target genes. BICORN prunes out genes regulated by less than two TFs or missing gene expression data. Then, based on the prior bindings on remaining genes, it automatically identifies a candidate CRM matrix including
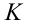
 rows and
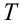
 columns. Each row vector represents a candidate CRM and each column denotes a particular TF.

Regarding candidate CRM searching, we extract all unique TF clusters from the prior binding pattern on existing genes. For each cluster, we count the number of genes regulated by this cluster. A parameter “*Minimum_gene_per_module_regulate*” is provided in the BICORN package to select TF clusters frequently showing up in the prior binding network. Then, a pool of candidate CRMs is identified. Assuming that there are
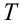
 TFs and
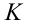
 CRMs, with
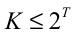
, the
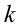
-th row in matrix
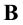
is a unique CRM vector
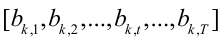
 with binary states
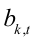
 as ‘0’ or ‘1’. In the real case, there are always some ‘background’ genes without any regulation. Hence, we include an all-zero vector
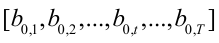
 in
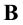
 to represent the ‘no regualtion’ case.

It is possible that a gene under regulation of multiple TFs (>2) in the prior binding network to be eventually regulated by a CRM with fewer TFs. For example, a gene regulated by three TFs in the prior binding network is possible to be regulated by only two TFs. If in the prior network a gene is regulated by 10 TFs, it can be eventually regulated by several CRMs with different TF members. Therefore, during the initialization process, when we assign candidate CRMs in the candidate CRM matrix
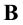
 to genes, we assign any candidate CRM containing a subset of prior TFs bindings to current gene.

Note that this step is achieved by the implementation of the R function “*data.initialization.R*”.

#### **S1.2 Gene expression data and log-linear modeling**

Gene expression data must be log-transformed and properly normalized. RNA-seq data with transcripts per kilobase million (TPM) values and microarray data both work. BICORN does not perform gene expression data normalization because it varies case by case and there exists many normalization tools performing this task. The input data are represented by a matrix (with
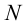
 genes as rows and
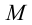
 samples as columns) of fold change of gene expression value of samples over the baseline expression. For time-course gene expression data, the input data should look like Time‘1’–Time‘0’, Time‘2’–Time‘0’, …, Time‘M’–Time‘0’, where expression under Time ‘0’ is referred to as baseline expression. For static gene expression data with multiple replicates measured under multiple conditions or with different treatments, the baseline expression can be obtained from control samples if there are any or the mean value across all samples. Then, the input data should look like Sample’1’-baseline, Sample’2’-baseline, …, Sample’M’-baseline.

We assume that the
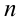
-th gene is regulated by the
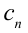
-th CRM. To infer
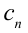
, gene expression data
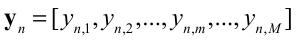
 are modeled using a log-linear equation as follows:

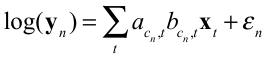
,	(S-1)

where vector
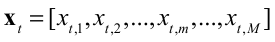
 represents the hidden TF activity (TFA) of the
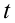
-th TF; variable
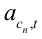
 represents the regulation strength of the
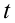
-th TF; and vector
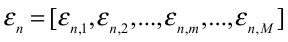
 denotes the fitting residue.

We denote matrices
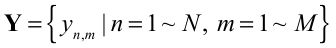
,
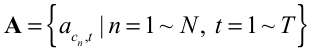
, and
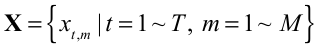
. To infer CRMs for all genes, we need to jointly estimate variables
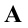
,
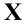
, and
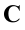
, and their joint posterior probability is defined as:

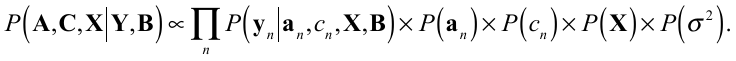
(S-2)

We make several key assumptions on the prior distribution of above variables. Specifically, the TFA vector
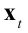
 is assumed to be a Gaussian random process and the prior distribution on each variable
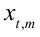
 is a Gaussian distribution
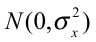
. Each regulation strength variable
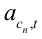
 is conditional on the state of
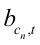
. For
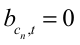
,
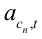
 equals to 0. For
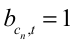
, we assume a prior Gaussian distribution on
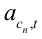
 as
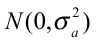
. The fitting residue variable is assumed to follow a zero-mean Gaussian distribution as
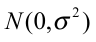
. The fitting residue variance
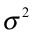
 is assumed to follow an inverse Gamma distribution as
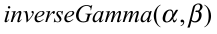
. We assume a uniform prior on candidate CRMs for each gene. Here,
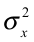
,
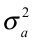
,
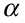
 and
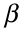
 are predefined hyper-parameters. Then, the above posterior probability can be further expanded as follows:

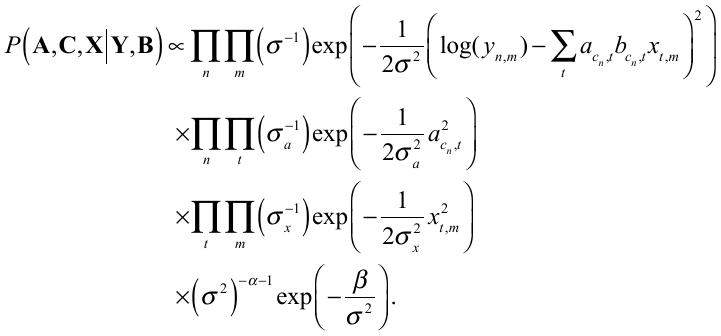
(S-3)

From Eq. (S-3), it is clear that the dependency exists on different variables. To estimate each variable, we iteratively calculate the conditional probability for each and perform estimation under a Gibbs sampling framework.

#### **S1.3 Gibbs sampling of TFA**

We sample TFA of the *t*-th TF under the *m*-th sample
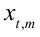
 according to its conditional probability as follows:

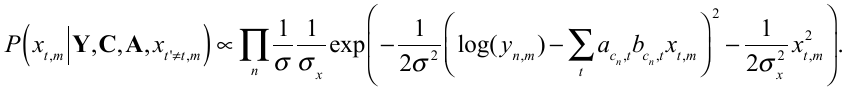
(S-4)

which is a Gaussian distribution with mean and variance parameters:

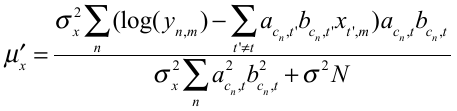
,
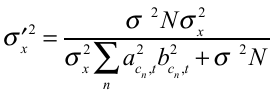
.	(S-5)

Note that we iteratively sample
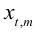
 because this probability is conditional on other TFAs
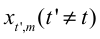
. For the hyper-parameter
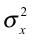
, we set an informative prior on
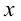
 with
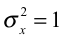
 since the gene expression data has been properly normalized.

Note that this step is achieved using the R function “***X_sampling.R***” in the package.

#### **S1.4 Gibbs sampling of regulation strength**

For each gene,
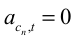
 if
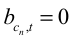
 (no binding); otherwise (if binding exists), we sample
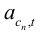
 according the following conditional probability:

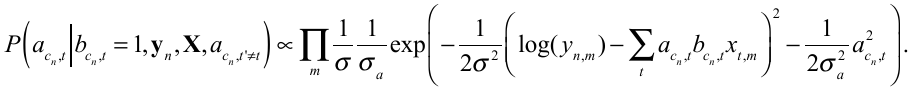
(S-6)

Specifically, the above conditional probability is a Gaussian distribution with mean and variance parameters as follows:

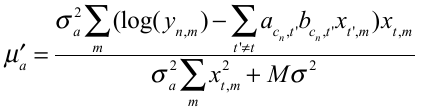
,
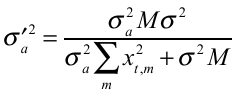
.	(S-7)

Note that we should iteratively sample
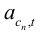
 for all TFs one by one because this probability is conditional on other
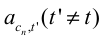
. In each round, we randomly shuffle the order of TFs to ensure that no bias is introduced. We assign non-informative large value to hyper-parameter
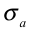
 as
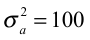
 to reflect that there is no assumed knowledge on regulation strength.

Note that in the BICORN package, this step is achieved using the R function “***A_sampling.R***”.

#### **S1.5 Gibbs sampling of CRM**

In order to sample a CRM for the *n*-th gene, we calculate a conditional probability for each CRM as follows:

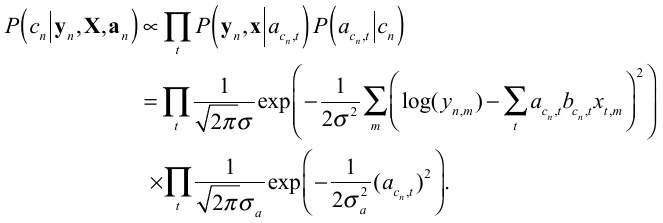
(S-8)

After calculating the above probabilities for individual CRMs, we sample
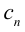
 by following probablily:
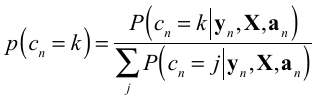
.

Note that this step is achieved using the R function “***C_sampling_cluster.R***”.

The overall fitting performance is controlled by the fitting residue variance
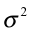
, which is further controlled using a prior inverse Gamma distribution. This variance should also be updated in each round of sampling to ensure the convergence of the overall sampling process. After estimating all above variables, we update
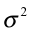
 according to its conditional probability:

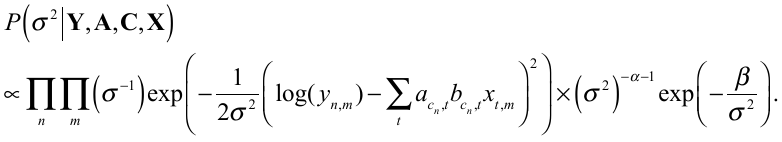
(S-9)

which is still an inverse Gamma distribution with updated parameters as follows:

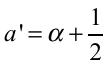
,
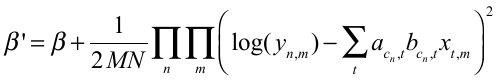
.	(S-10)

We set hyper-parameters
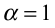
and
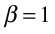
 to make the prior distribution of
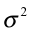
 non-informative.

In the BICORN R package, this step is achieved using the R function “***sigmanoise_sampling.R***”. The above-mentioned sampling steps are iteratively carried out in the main function “***BICORN.function.R***”

### **S2 Simulation study and performance comparison**

#### **S2.1 CRM-gene interaction reconstruction**

Existing integrative methods can only infer individual TF bindings rather than CRMs. For a target gene, if only partial bindings are inferred, the module or the association of multiple TFs is not correctly or fully reconstructed. Using our BICORN R package and two existing R software packages: BNCA (Sabatti and James, 2006) and COGRIM (Guan, et al., 2014), we evaluate whether the CRM of a gene can be completely reconstructed. We simulated a prior binding network including 160 genes and 20 TFs. There are 80 true target genes, and each is regulated by a CRM with 2 ~ 6 TFs. Regulation strengths were generated by following a Gaussian distribution. TFAs for individual TFs were simulated using a Gaussian random process with zero mean and standard deviation under 20 samples. Then, we simulated a gene expression dataset using log-linear model as in Eq. (S-1).

We use two performance metrics, precision and recall, to evaluation how well the CRMs of genes can be reconstructed; precision and recall are defined by:

precision = (Number of genes with correct CRMs)/(Number of genes identified)

recall = (Number of genes with correct CRMs)/(The total number of true target genes)

**Table S1.** Precision and Recall performance on CRM inference for target genes

| **METHOD** | **BICORN** | **BNCA** | **COGRIM** |
| --- | --- | --- | --- |
| **Precision** | 0.768 | 0.730 | 0.725 |
| **Recall** | **0.663** | 0.100 | 0.090 |

It can be seen from Table S1 that to achieve the same precision performance, BICORN identified CRMs for a significantly larger number of true target genes than BNCA or COGRIM, with a recall value 0.66 vs. ~0.1. Evidently, conventional regulatory network inference tools like BNCA or COGRIM cannot recover CRMs effectively.

#### **S2.2 TF-gene network reconstruction**

Using simulation data, we can further demonstrate that for conventional TF-gene binding inference, BICORN can provide an improved performance. We simulated different noise scenarios by varying the false positive rate of initial binding connections (from 0.05 to 0.25 with step 0.05) and the signal-to-noise ratio (SNR) of gene expression data (from 9 dB to -3 dB with a step of 3 dB). We included two more regulatory network inference tools, NARROMI (Zhang, et al., 2013) and a LASSO based approach (Qin, et al., 2014), for comparison.

We calculate the sensitivity and specificity as follows:

,

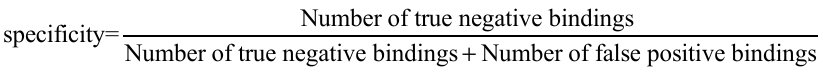
.

Area under receiver operating characteristic (ROC) curves (AUCs) of BICORN and competing methods are calculated from the sensitivity and specificity, which are shown in Fig. S2.

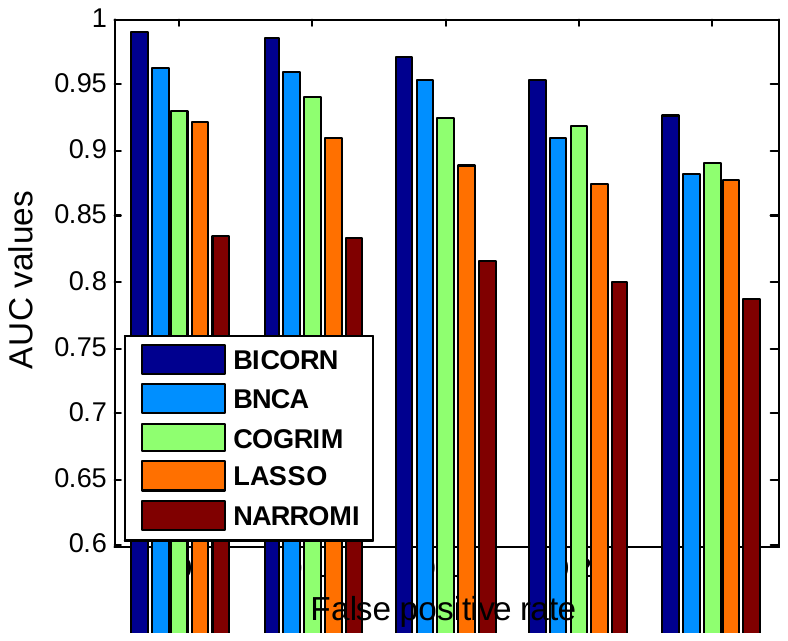

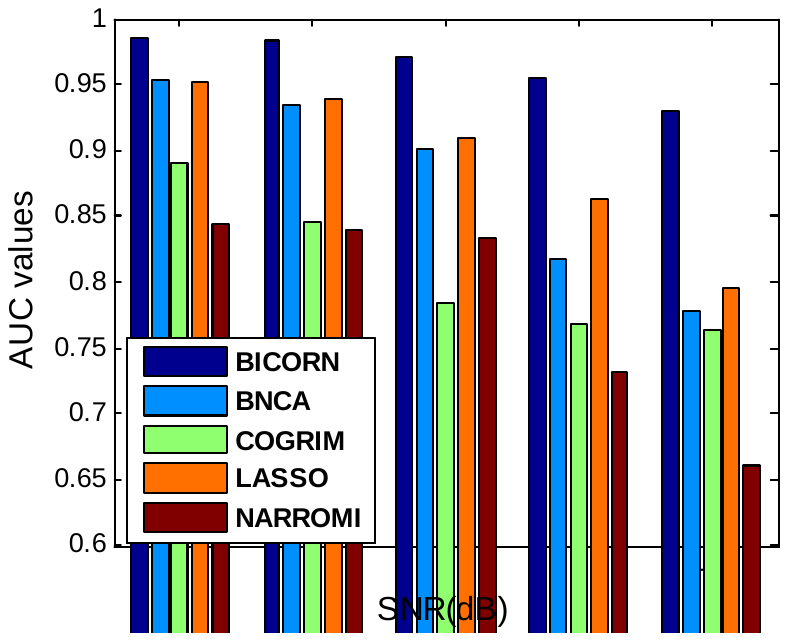

(A)	(B)

**Figure S2.** Binding network prediction performance of competing methods: (A) initial binding networks with different false positive rates; (B) gene expression data with different Signal-to-Noise Ratio (SNR).

#### **S2.3 Target gene identification**

In biological studies, researchers are interested in identifying functional target genes, especially given a large number of TFs. COGRIM was a main method previously proposed to identify target genes given gene expression data and a noisy prior TF-gene binding network. We simulated several more challenging scenarios with noisy prior bindings of high false positive rate. For each method, a true target gene is called if at least one binding with a sampling frequency over a default threshold. In this case, we calculate the sensitivity and specificity as follows:

,

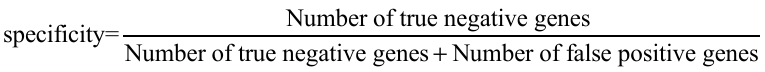
.

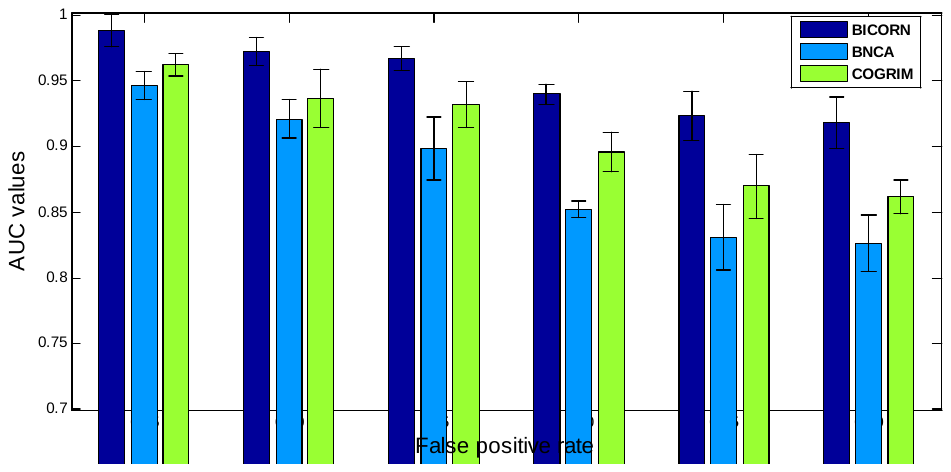

**Figure S3.** AUC performances of competing methods on target gene prediction using noisy physical binding networks.

As shown in Fig. S3, BICORN is quite robust in identifying target genes from data with false negatives. This robust performance can be mainly attributed to that the BICORN algorithm is specifically designed to act on modules (i.e., CRMs) directly. Since a background gene is not truly regulated by any TFs, there is no TF combination showing a significantly higher probability than the others. For such genes, BICORN reports all ‘0’ bindings. For competing methods, individual TF bindings are evaluated for each gene so that the chance of inferring a false positive binding or a target gene is higher, leading to a degrading performance on identifying true target genes.

### **S3 *De novo* CRM inference using breast cancer MCF-7 data**

We applied BICORN to real data acquired from breast cancer MCF-7 cells. We downloaded peak files of 39 TFs from ENCODE (https://www.encodeproject.org/). To infer MCF-7 cell-specific CRMs, we downloaded two 17b-E2 treated MCF-7 RNA-seq data sets from the GEO database (accession numbers GSE62789 and GSE51403). It is not a trivial task to directly validate CRMs. To test the robustness of BICORN on CRM inference, we applied BICORN to the same prior binding knowledge but with two relevant gene expression datasets (one time-course dataset and one static dataset with similar treatments). We processed both datasets using RSEM for gene expression measurements (Li and Dewey, 2011). The GSE62789 dataset includes 10 RNA-seq samples measured at baseline (‘0’ time point) and 9 different time points within 24 hours (hrs) of 10nM 17b-E2 treatment. The GSE51403 dataset includes 7 RNA-seq samples generated under vehicle condition and another 7 RNA-seq samples generated after 24hrs treatment of 10nM 17b-E2. In total, we collected differentially expressed genes as reported in the original publications and identified 275 common target genes for further exploration.

Gene promoter region is defined as +/1k bps from the nearest transcription start site. We annotated TF peaks using GREAT (McLean, et al., 2010). Based on the prior binding events (annotated TF peaks) on 275 target genes, BICORN identified 73 candidate CRMs. After 1,000 rounds of Gibbs sampling, BICORN inferred 549 reliable CRM-gene interactions ((sampling frequency > 0.85)) using the time-course gene expression dataset and 545 reliable CRM-gene interactions using the steady-state gene expression dataset. As shown in Fig. S4(A), in total there are 466 common CRM-gene interactions, with an overlap of 86%. Based on the number of target genes regulated by each CRM, the top-ranked 20 CRMs are presented in Fig. S4(B).

We also downloaded MCF-7-specific enhancer-like regions and ChIA-PET data from the ENCODE database. BICORN identified 56 candidate CRMs after examining binary binding events at enhancers associated with above 275 target genes. For each target gene, BICORN will evaluate the regulatory performance of candidate CRMs from all mapped enhancers and identify the final CRM(s) distantly regulating the expression of current gene. After 1,000 rounds of Gibbs sampling, BICORN identified 822 reliable CRM-gene interactions from the time-course gene expression dataset and 816 reliable CRM-gene interactions from the static gene expression dataset. As shown in Fig. S5(A), in total there are 630 common CRM-gene interactions, with an overlap of 77%. Based on the number of target genes regulated by each module, the top-ranked 20 CRMs are presented in Fig. S5(B).

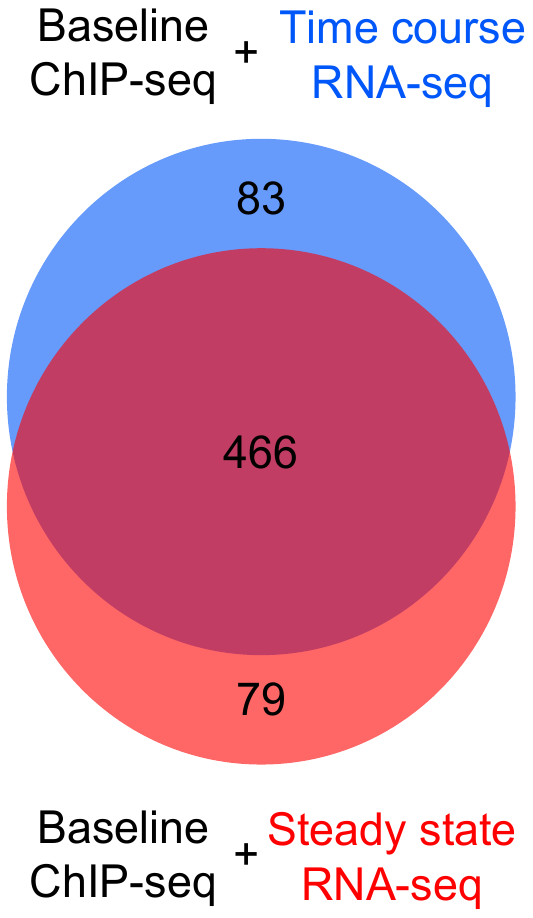

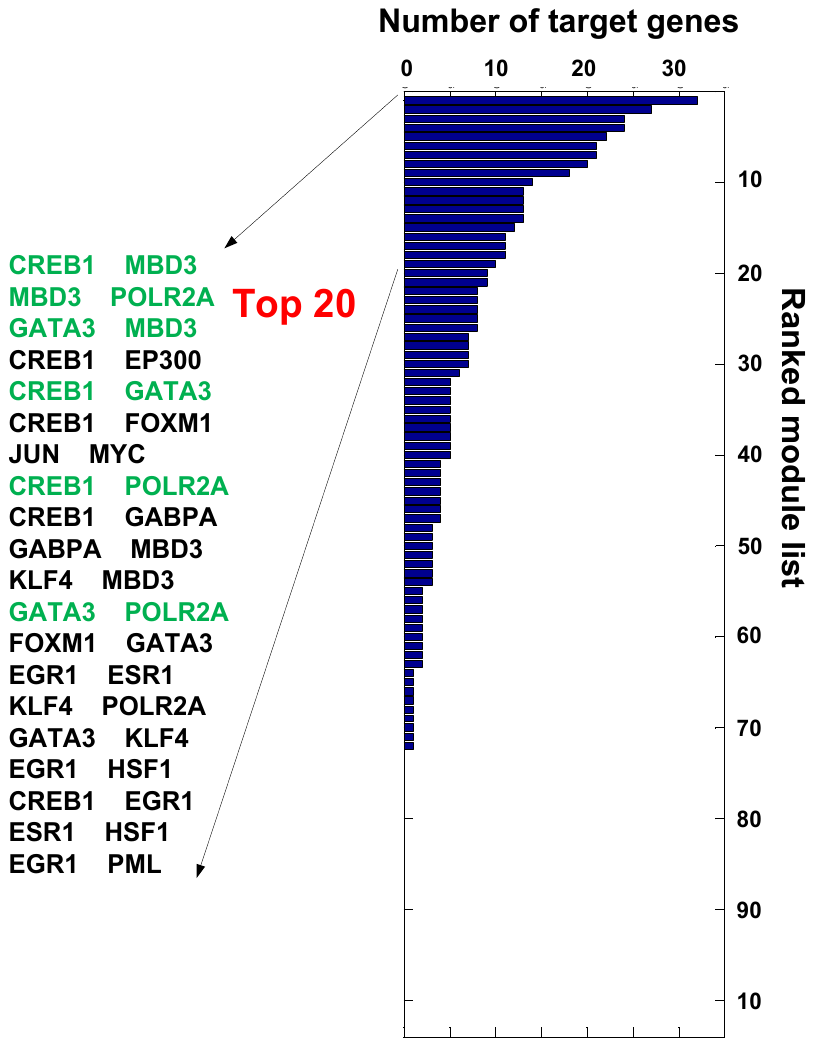

(A)	(B)

**Figure S4.** BICORN integrative analyses of 32 TFs at gene promoter regions: (A) Venn diagram of BICORN inferred functional bindings from two different gene expression data sets; (B) the top 20 of 73 BICORN-identified CRMs, sorted by the number of active target genes.

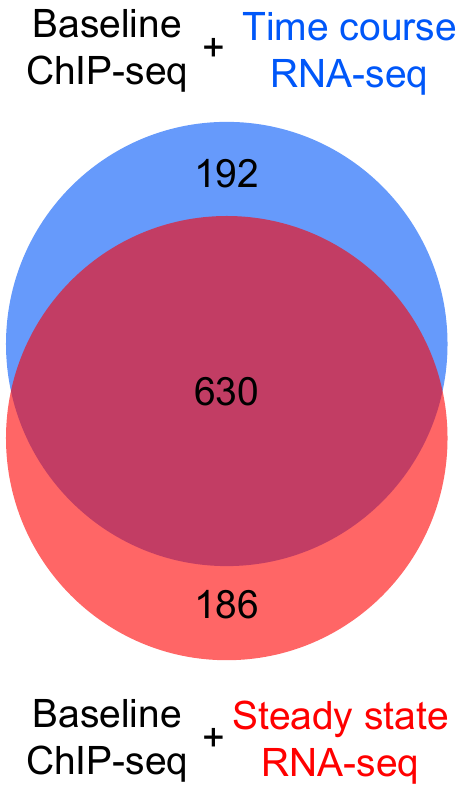

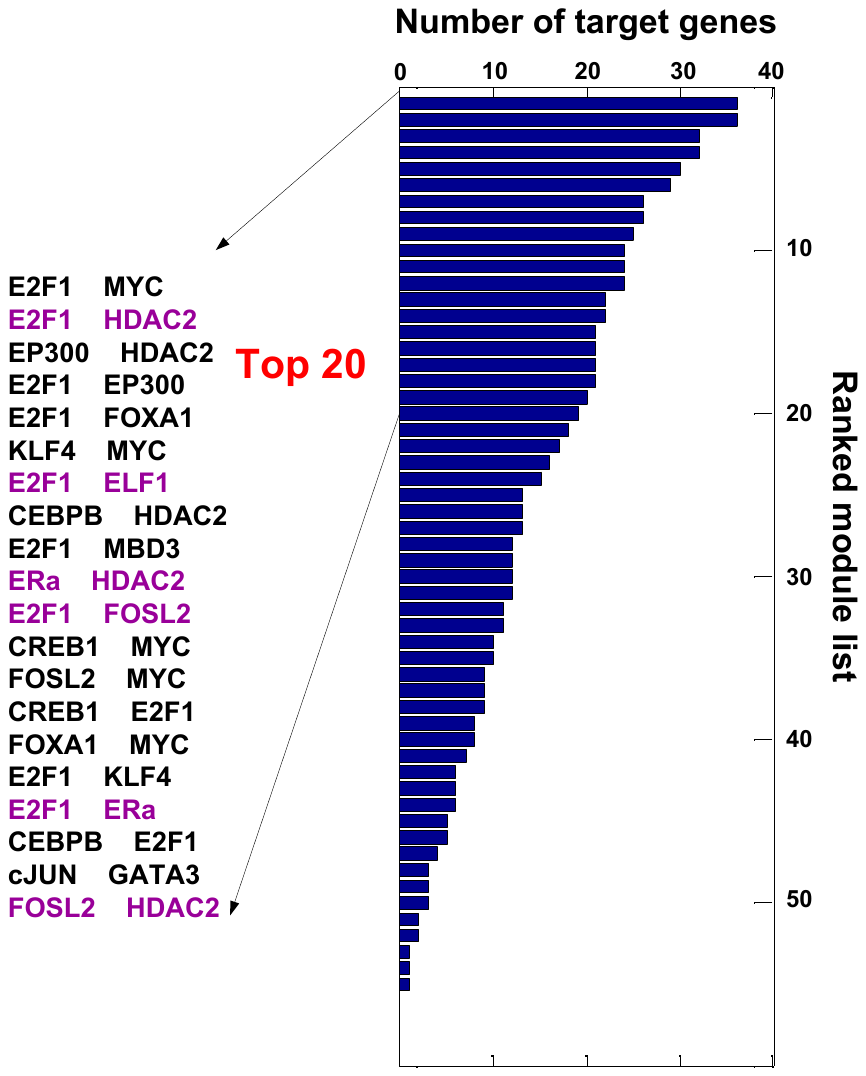

(A)	(B)

**Figure S5.** BICORN integrative analyses of 22 TFs at enhancer regions: (A) Venn diagram of BICORN inferred functional bindings from two different gene expression data sets; (B) the top 20 of 56 BICORN-identified CRMs, sorted by the number of active target genes.

Comparing the above two studies specific for breast cancer MCF-7 cells, we found that active TFs at promoter and enhancer regions are very different, as illustrated in Fig. S6. There are 7 TFs including HSF1, cJUN, EGR1, GABPA, FOXM1, MBD3 and GATA3 with intensive bindings at gene promoter regions, as shown in Fig. S6(A). However, those TFs have no strong functional bindings at enhancer regions, as can be seen from Fig. S6(B). Conversely, there are 6 TFs including E2F1, FOSL2, ELF1, HDAC2, CEBPB and FOXA1 highly active at enhancer regions. Their bindings at promoter regions are non-functional. Evidently, regulatory mechanisms at promoter and enhancer are quite different. Therefore, we recommend that CRMs should be inferred for TF binding events at promoter and enhancer regions, respectively.

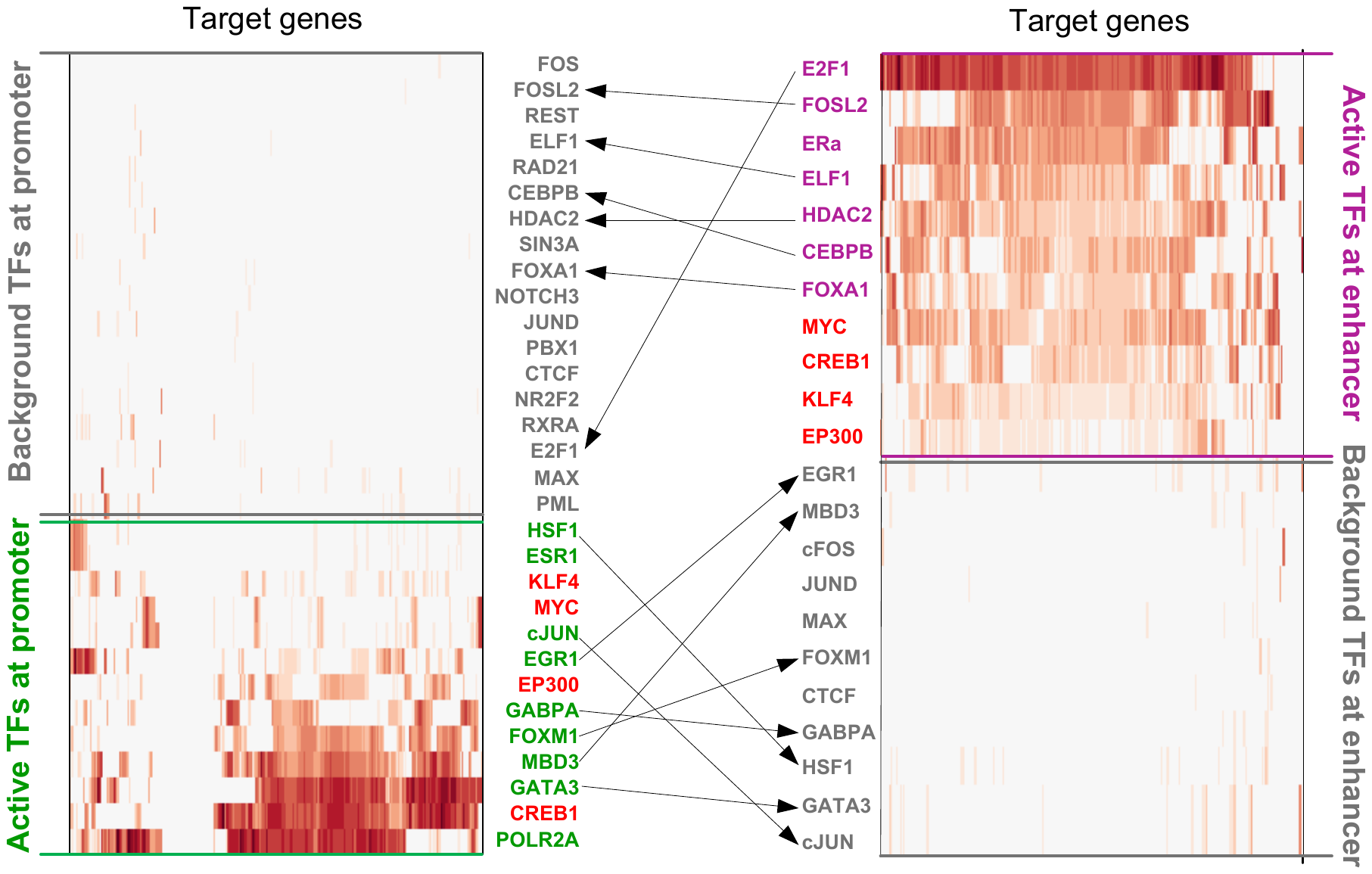

(A)	(B)

**Figure S6.** TFs functional at promoter or enhancer regions of E2 responsive target genes in breast cancer MCF-7 cells: (A) foreground (green) and background (grey) TFs at promoter regions; (B) foreground (purple) and background (grey) TFs at enhancer regions. Common foreground TFs at both types of regions are labeled as ‘red’.

### **S4 *De novo* CRM inference in different cell types**

To demonstrate the broad applicability of BICORN, 6 cell types were selected each with ChIP-seq data of at least 20 TFs available in the ENCODE database. For each cell type, a matched gene expression data set was downloaded from the GEO database, as summarized in Table S2. For CRM inference at enhancer regions, the selection of a cell type depends on whether there is a collection of cell type-specific enhancer-like regions in the ENCODE database.

**Table S2** Prior bindings and gene expression data used for CRM inference

| **Cell line** | **Number of TFs (ENCODE database)** | **Gene expression data (GEO database)** | **Promoter** | **Enhancer** |
| --- | --- | --- | --- | --- |
| K562 | 203 | GSE1036 | √ | √ |
| GM12878 | 122 | GSE51709 | √ | √ |
| HepG2 | 108 | GSE6869 | √ | √ |
| A549 | 52 | GSE69667 | √ |  |
| SK-N-SH | 28 | GSE9169 | √ |  |
| HCT116 | 20 | GSE14103 | √ | √ |

***K562***

A K562 microarray data set was downloaded from the GEO database (GSE1036). Expression data were analyzed by MAS 5.0. According to (Addya, et al., 2004), 1,311 differentially expressed genes were selected. We downloaded K562-specific ChIP-seq peaks of 203 TFs. For the promoter study, we overlapped above peaks with the promoter region of differential genes and identified 68 candidate CRMs for further exploration. After 1000 rounds of sampling, candidate CRMs were prioritized by the number of regulated target genes (posterior probability threshold for confident CRM-gene regulations is 0.9), a set of top-ranked CRMs (21 CRMs, each regulating at least 10% of the original set of target genes) were identified. Among them, one example of the high ranked CRMs is CTCF-RAD21. This module has been previously reported to be functional around transcription starting sites of genes in K562 cells. ChIP experiments showed that enrichment of CTCF was significantly reduced (average of threefold) in K562 cells subjected to Rad21 knock-down as compared to control K562 cells, suggesting that RAD21 contributes to stable CTCF bindings (Hou, et al., 2010).

For the enhancer study, we downloaded K562-specific enhancer-like regions and ChIA-PET data from the ENCODE database. We overlapped TF peaks with enhancer regions that can be mapped to differential genes through enhancer-promoter interactions (each interaction with at least two paired-end ChIA-PET reads). In total, we collected 2,135 enhancers regions, 607 genes and 82 candidate CRMs for further exploration. Among the BICORN-identified CRMs (20 CRMs), NRF1-RFX1 is the top-ranked CRM. RFX1 is annotated in Gene Ontology as “*RNA polymerase II distal enhancer sequence-specific binding”.*

***GM12878***

A GM12878 microarray data set was downloaded from the GEO database (GSE51709). Expression data were analyzed through Affymetrix Expression Console. According to (Su, et al., 2015), 696 differentially expressed genes were selected. We downloaded GM12878 specific ChIP-seq peaks of 122 TFs the ENCODE database. For promoter study, we overlapped above peaks with the promoter region of differential genes and identified 79 candidate CRMs for further exploration. After 1000 rounds of sampling, 17 confident CRMs were identified. As an example, the top-ranked CRM is ELF1-ZEB1; the appearance of both in this GM12878 (B-Lymphocyte in blood tissue) study is highly expected since both of them are B-cell specific transcription factors.

For the enhancer study, we downloaded GM12878-specific enhancer-like regions and ChIA-PET data from the ENCODE database. We overlapped TF peaks with enhancer regions which can be mapped to differential genes through enhancer-promoter interactions (each interaction with at least two paired-end ChIA-PET reads). In total, we collected 1571 enhancers regions, 381 genes and 68 candidate CRMs for further exploration. After 1000 rounds of sampling, 10 confident CRMs were identified. The top-ranked CRM is MTA2-TBX21; MTA2 occupancy on enhancers was previously demonstrated in (Zhang, et al., 2015).

***HepG2***

A HepG2 microarray data set was downloaded from the GEO database (GSE6869). Expression data were obtained by MAS 5.0. According to (De, et al., 2010), 874 differentially expressed genes were selected. We downloaded HepG2 specific ChIP-seq peaks of 108 TFs the ENCODE database. For the promoter study, we overlapped above peaks with the promoter region of differential genes and identified 55 candidate CRMs for further exploration. After 1000 rounds of sampling, 12 confident CRM were identified. The top-ranked CRM is SOX13-SOX5; both transcription factors are from SOX family and it has been known that SOX13 complements SOX5 functionally (Baroti, et al., 2016).

For the enhancer study, we downloaded HepG2-specific enhancer-like regions and ChIA-PET data from the ENCODE database. We overlapped TF peaks with enhancer regions which can be mapped to differential genes through enhancer-promoter interactions (each interaction with at least two paired-end ChIA-PET reads). In total, we collected 1457 enhancers regions, 433 genes and 43 candidate CRMs for further exploration. After 1000 rounds of sampling, 8 confident CRMs were identified. One of the top-ranked CRMs has two members as SUZ12 and ZNF143, two PRC2 epigenomic signatures, whose associations with enhancers have been reported (Dozmorov, 2015).

***A549***

An A549 RNA-seq data set was downloaded from the GEO database (GSE69667). Expression data was analyzed through RSEM. According to (Chang, et al., 2016), 1,633 differentially expressed genes were selected. We downloaded A549 specific ChIP-seq peaks of 52 TFs from the ENCODE database. Due to lack of A549 specific enhancer-like regions in the ENCODE database, in this case we conducted CRM inference at gene promoter region only. We overlapped ChIP-seq peaks with the promoter region of selected 1,633 genes and finally identified 72 candidate CRMs for further exploration. After 1000 rounds of sampling, 13 confident CRMs were identified. An example of the top-ranked CRMs is E2F6-MAX; MAX is a member of the E2F6 complex that usually binds to E2F-reposive promoters and it is in involved in gene repression (Ogawa, et al., 2002).

***SK-N-SH***

A SK-N-SH microarray data set was downloaded from the GEO database (GSE9169). Expression data was analyzed through Affymetrix Expression Console. According to (Nishida, et al., 2008), 362 differentially expressed genes were selected. We downloaded SK-N-SH specific ChIP-seq peaks of 52 TFs from the ENCODE database. Due to lack of SK-N-SH specific enhancer-like regions in the ENCODE database, in this case we conducted CRM inference at gene promoter region only. We overlapped ChIP-seq peaks with the promoter region of selected genes and finally identified 61 candidate CRMs for further exploration. After 1000 rounds of sampling, 14 confident CRMs were identified. One of the top-ranked CRMs includes three transcription factors as CTCF, RAD21 and SMC3. CTCF and the cohesin complex, consisting of the core subunits as SMC3 and RAD21 were found to colocalize extensively throughout mammalian genomes (Wendt, et al., 2008).

***HCT116***

A HCT116 microarray data set was downloaded from the GEO database (GSE14103). Expression data was analyzed through Affymetrix Expression Console. According to (Mizuno, et al., 2009), 286 differentially expressed genes were selected. We downloaded HCT116 specific ChIP-seq peaks of 20 TFs from the ENCODE database. For the promoter study, we overlapped above peaks with the promoter region of differential genes and identified 55 candidate CRMs for further exploration. After 1000 rounds of sampling, 24 confident CRMs were identified. Among them, the top-ranked CRM is FOSL1-JUND. It is not a rare case to find this module since FOSL1 requires a dimerization partner to regulate gene transcription and this partner is often a member of the JUN family (Shaulian and Karin, 2002).

For the enhancer study, we downloaded HCT116-specific enhancer-like regions and ChIA-PET data from the ENCODE database. We overlapped TF peaks with enhancer regions which can be mapped to differential genes through enhancer-promoter interactions (each interaction with at least two paired-end ChIA-PET reads). In total, we collected 366 enhancers regions, 110 genes and 44 candidate CRMs for further exploration. After 1000 rounds of sampling, 6 confident CRMs were identified. An example of the top-ranked CRMs is POLR2A-YY1, which generally occupies active enhancers and promoters across cell types and play structural roles in enhancer-promoter loops (Weintraub, et al., 2017).
